# Synaptic and intrinsic plasticity coordinate spike output in cerebellar Purkinje cells

**DOI:** 10.1101/742577

**Authors:** Hyun Geun Shim, Sang Jeong Kim

## Abstract

Learning has been thought to be implemented by activity-dependent modifications of synaptic weight and intrinsic excitability. Here, we highlight how long-term depression at parallel fiber to Purkinje cell synapses (PF-PC LTD) and intrinsic plasticity of PCs coordinate the postsynaptic spike discharge from C57BL/6 male mice. Intrinsic plasticity of PCs in the flocculus matched the timing rules and shared intracellular signaling for PF-PC LTD. Notably, the intrinsic plasticity was confined to the dendritic branches where the synaptic plasticity is formed. Besides, when either synaptic or intrinsic plasticity was impaired, the impact of PF inputs was less reflected by the spike output of PCs. In conclusion, synergies between synaptic and intrinsic plasticity may play a role in tuning the PC output, thereby achieving optimal ranges of output.

## Introduction

Emerging evidence has revealed that learning and memory is implemented not only by activity-dependent synaptic plasticity but also by non-synaptic intrinsic plasticity in several neural circuits (Crestani et al., 2018; Lisman et al., 2018; Shim et al., 2018; Zhang and Linden, 2003). Of interest, intrinsic excitability contributes to integration of synaptic inputs and generation of net neuronal output (Hoffman et al., 1997; Lev-Ram et al., 2003, 1995; Shim et al., 2017). Therefore, the plasticity of intrinsic excitability may incorporate synaptic plasticity, and thereby formation of postsynaptic spike output (Najac and Raman, 2015). In spite of the physiological significance, the precise role of how intrinsic plasticity interacts with synaptic plasticity to shape information processing remains elusive.

Cerebellar PC intrinsic plasticity has been implicated as a neural computational feature of the information processing of cerebellar cortex (Belmeguenai et al., 2010; Grassi and Pettorossi, 2001; Shim et al., 2018, 2017; Steuber et al., 2007). Notably, bidirectional modulation of synaptic plasticity at the PF-PC synapses has been found to be accompanied by intrinsic plasticity, which shows the same polarity with synaptic plasticity, indicating that the intrinsic plasticity might synergistically enable the PC spike output to be sufficiently modulated when synaptic plasticity occurs (Belmeguenai et al., 2010; Shim et al., 2017). Furthermore, multiple lines of evidence have suggested that non-synaptic intrinsic plasticity of PCs might be another player involved in information storage for cerebellar motor learning (Jang et al., 2019; Ryu et al., 2017; Schonewille et al., 2011). However, how intrinsic plasticity contributes to input-output coordination in accordance with learning is still unclear.

In this work, linking of PF-PC LTD and LTD of intrinsic excitability (LTD-IE) was found to be indispensable for robust reduction of PC spiking output after formation of cerebellar LTD. Inhibition of either PF-PC LTD or LTD-IE itself affected PC spike probability, however, prominent changes in spike output were produced by the synergies between both forms of PC plasticity. Interestingly, the effects of synergy on PC output signals were found to be spatially restricted within the conditioned dendritic branch where synaptic LTD is formed. Thus, the synaptic input through the unconditioned dendritic branches have less impact on PC spike output.

## Results and Discussion

### Timing rules of intrinsic plasticity of floccular PCs

All of the electrophysiological whole-cell patch clamp recordings were executed from cerebellar flocculus slices obtained from 4 – 6 weeks old male C57BL/6 mice (figure 1A). In the previous study, the PF-PC LTD in the flocculus, sub-region of the cerebellum supporting a oculomotor learning, was found to require distinct timing rules of PF and CF activation from that in the spinocerebellum (lobule III to V). We first tested whether intrinsic plasticity of PCs in the flocculus follows the timing rule for governing the PF-PC LTD by introducing three previously verified protocols for LTD induction: simultaneous stimulation of PF and climbing fiber (CF) (PF-LTD_ISI=0_), stimulation of PF and CF with 120 ms of intervals (PF-LTD_ISI=120_) and 100 Hz of burst stimuli of PF followed by stimulation of CF with 150 ms intervals (PF-LTD_ISI=150burst_) (figure 1B and see Material and Method) (Shim et al., 2017; Suvrathan et al., 2016). LTD-IE was produced by the LTD-inducing protocols, whereas PF-LTD_ISI=0_ failed to induce the LTD-IE (figure 1C – F). In the following experiments, the PF-LTD_ISI=120_ was used to induce PF-PC LTD and LTD-IE. In addition, the LTD-IE in the floccular PCs was dependent on the activation of metabotropic glutamate receptor 1 (mGluR1) and Ca^2+^/calmodulin-dependent protein kinase II (CaMKII), implying that both forms of plasticity in PCs may share the same intracellular signaling cascade (Supplemental figure 1). We further investigated the impact of synaptic plasticity on postsynaptic spiking output. PC spike output was measured by counting the number of spikes elicited by PF synaptic stimulation (20 PF stimuli at 20Hz, 1s). Compared to baseline, the PF-evoked spike output robustly decreased after PF-PC LTD induction when the PF-PC LTD was concomitant with LTD-IE, without changes in EPSP summation (figure 1G – I). On the other hand, when LTD and LTD-IE were not induced, there was no significant change of PF-evoked spike count.

**Figure 1.**
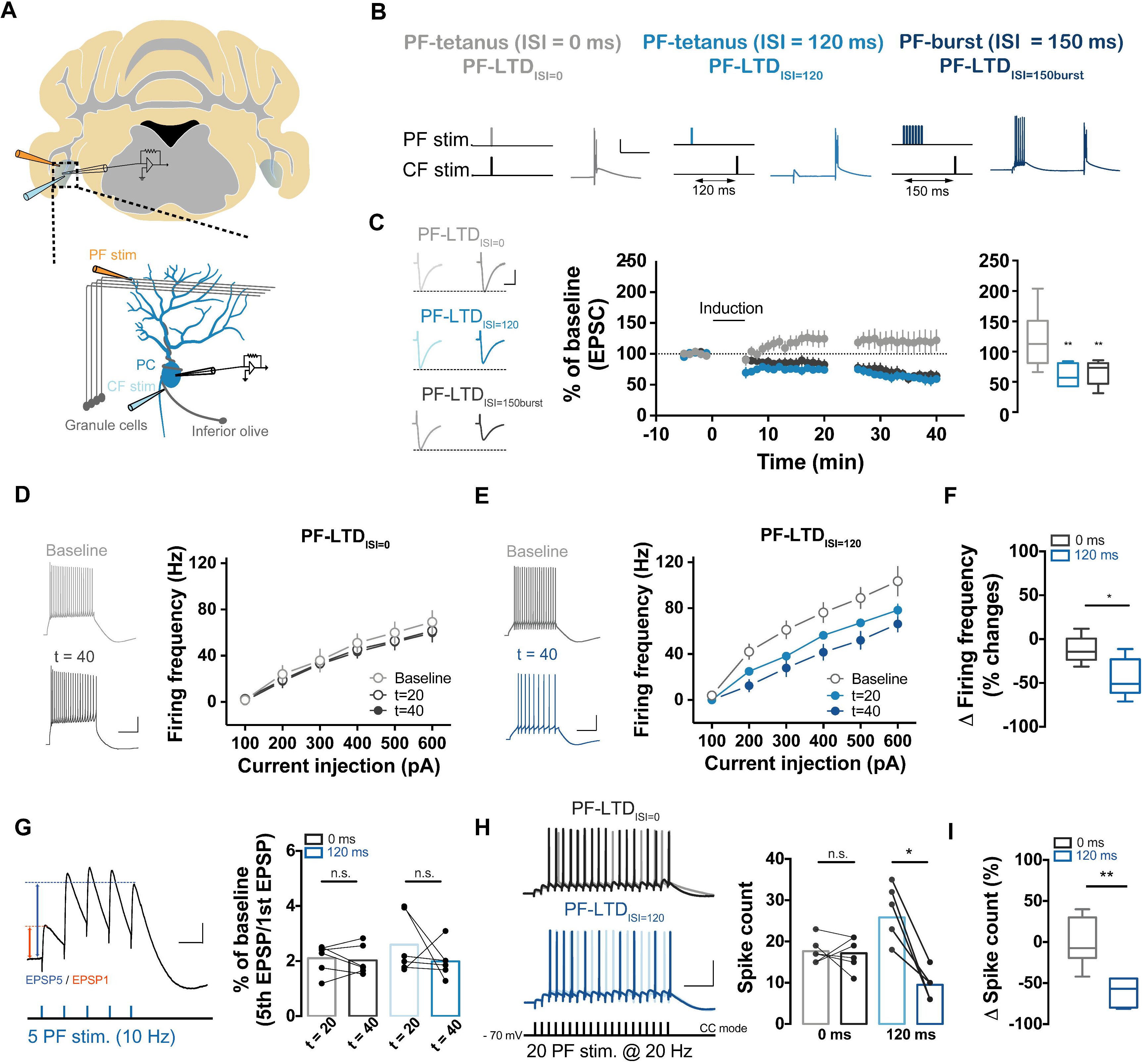
Shared Timing rules for induction of PF-PC LTD and LTD-IE. (A) Illustration of the cerebellar flocculus (upper) and the recording site for synaptic plasticity (bottom). (B) Three protocols were used for induction of LTD and LTD-IE. Tetanizing of PF with 1 Hz for 5 min was delivered and single CF was stimulated simultaneously or following with inter-stimulation interval (ISI) of 120 ms [PF-LTD_ISI=0_, left (grey) vs. PF-LTD_ISI=120_, middle (blue)], or the PF-burst protocol consisted of 7 times of burst stimulation onto PF with 100 Hz followed by CF stimulation with ISI of 150 ms in every 10 s for 5 min [PF-LTD_ISI=150burst_), right (dark blue)]. (C) Plots and summarizing box and whisker plots (middle and right) of changes in eEPSC from PF-LTD_ISI=0_ (grey, n = 7), PF-LTD_ISI=120_ (blue, n = 7) and PF-LTD_ISI=150burst_ (dark blue, n = 9). Consistent to previous observation, delayed activation of CF with 120 ms and 150 ms of delay from PF activation induced PF-LTD (F = 9.256, p = 0.001, One-way ANOVA). Insets (left) show the representative trace of eEPSC before and after induction. Scale bar: 25 ms (horizontal) and 50 ms (vertical). (D - E) Plots showing frequency – current (F/I) curve of PF-LTD_ISI=0_ (D: black; n = 5) and PF-LTD_ISI=120_ (E: blue; n = 5). Insets show representative traces of depolarization-induced AP train. Intrinsic excitability was significantly decreased after LTD induction with PF-LTD_ISI=120_ (p = 0.03, Mann-Whitney test). Scale: 200 ms (horizontal) and 20 mV (vertical). (F) Summarizing box and whisker plots showing comparison of ∆ firing frequency between groups (∆firing frequency: PF-LTD_ISI=0_ = −12.06 ± 6.95%, n = 5; PF-LTD_ISI=120_ = −43.87 ± 9.98%, n = 5, p = 0.03, Mann-Whitney). (G) Bar graph showing the changes in temporal summation of the EPSP from two protocols (black: PF-LTD_ISI=0_; blue: PF-LTD_ISI=120_). Summation was determined by calculating the ratio of 5th EPSP amplitude to 1st EPSP amplitude (PF-LTD_ISI=0_: p = 0.71, n = 6; PF-LTD_ISI=120_: p = 0.36, n = 6). Insets (right) show a representative trace and protocol. EPSP summation was not changed after LTD induction. Scale: 100 ms (horizontal) and 5 mV (vertical). (H) Bar graphs showing the changes in PF-evoked spike count between before and after induction from the groups (grey: PF-LTD_ISI=0_ before induction; black: PF-LTD_ISI=0_ after induction, n = 6; light blue: PF-LTD_ISI=120_ before induction; blue: PF-LTD_ISI=120_ after induction, n = 7). Only in PF-LTD_ISI=120_ showed the significant reduction of the PF-evoked spike count (PF-evoked spike count: baseline = 25.83 ± 2.96 vs. t = 40 = 9.50 ± 1.36, p = 0.03, Wilcoxon test) compared to PF-LTD_ISI=0_ (PF-evoked spike count: baseline = 17.67 ± 1.23 vs. t = 40 = 17.17 ± 1.60, p = 0.81, Wilcoxon test). Insets show representative traces of PF-evoked spikes, elicited by stimulating 20 times of PF with 20 Hz. Scale: 250 ms (horizontal) and 20 mV (vertical). (I) Box and whisker plots showing the PF-evoked spike count from PF-LTD_ISI=0_ (grey) and PF-LTD_ISI=120_ (blue). LTD-inducing protocol robustly decreased the PF-evoked spiking activity (∆spike count: PF-LTD_ISI=0_ = 0.30 ± 12.02% vs. PF-LTD_ISI=120_ = −60.52 ± 6.65%, p = 0.002, Mann-Whitney test). For statistics, one-way ANOVA was used for C (righy panel) and post hoc Tukey’s test was used for comparison between groups. Two-way RM ANOVA was used for D and E and post hoc Tukey’s test was used for within in group (time) comparison. Mann-Whitney test was used for F and I. Wilcoxon test was used for comparison of paired data set in G and H. Error bar indicates SEM. n.s. denotes ‘not significant’; *P < 0.05, **p < 0.01.

### Conditioned PF branches contribute to robust reduction of spike output of the PCs

Several lines of evidence have described that each individual dendritic branch of a neuron could be an information processing unit (Belmeguenai et al., 2010; Zang et al., 2018). Thus, we tested if the input-ouput coordination would be tuned within a specific dendritic branch after formation of synaptic LTD and LTD-IE by comparing spike probability in response to electrical PF stimulation delivered onto the two different sites of PF beams (figure 2A and B). During the induction period, the PF-PC LTD protocol was delivered at the conditioning site (conditioned PF) whereas the PF tetanizing was omitted at the other branch site (unconditioned PF). PF-PC LTD was produced at only the dendritic branches where the tetanus stimuli were delivered and excitability change was also shown after LTD induction as shown in figure 1 (figure 2C and D), indicating the local-specificity of the synaptic plasticity. Interestingly, the PF-evoked spike count robustly decreased when the conditioned PF was stimulated. On the other hand, unconditioned PF-evoked spike count showed a slight reduction compared to baseline (−52.28 ± 4.35% vs. −11.71 ± 5.97%, conditioned and unconditioned PF, respectively, p < 0.001, Mann-Whitney test; figure E and F). Thus, synaptic LTD and LTD-IE were confined to the specific branches, thereby synergistically coordinating input-output relationship

**Figure 2.**
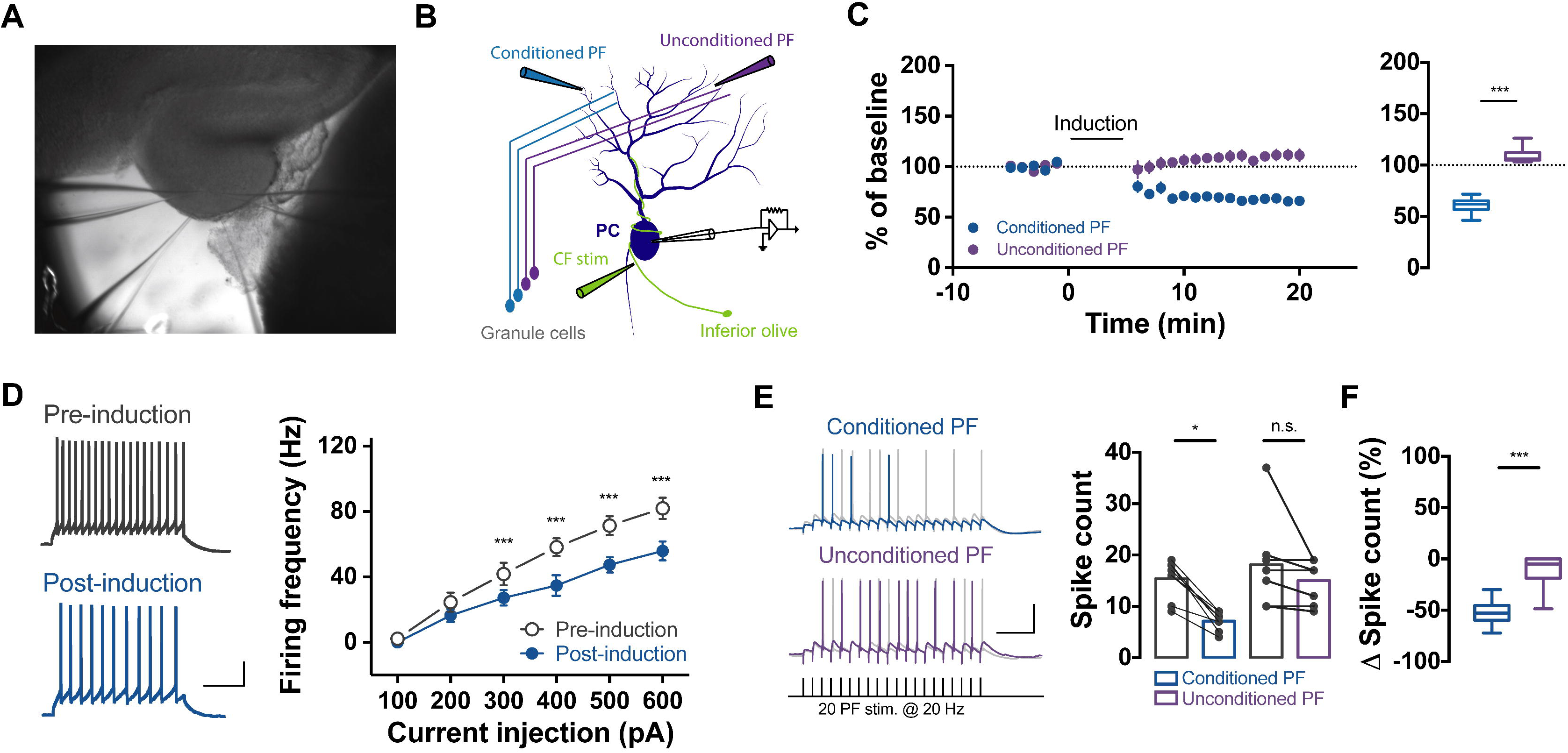
Synergistic plasticity of the PC spike output in a dendritic branch-specific manner. (A - B) DIC image and illustration of experimental strategy. Two sites of PF branches of the one neuron were stimulated. (B) LTD-inducing protocol was delivered onto the one site of PF branch (conditioned PF, blue) and tetanizing was omitted in the other side of PF branch (unconditioned PF, purple). (C) Plots (left) showing the normalized eEPSC before and after LTD induction in the two different branches and summarizing box and whisker plots (right) of changes in eEPSC of conditioned PF (blue, n = 8) and unconditioned PF (purple, n = 8). PF-PC LTD occurred only in the conditioned PF (% of baseline: conditioned PF = 67.16 ± 3.51% vs. unconditioned PF = 110.10 ± 5.05%, Mann-whitney test). (D) Plots showing frequency – current (F/I) curve pre and post induction of LTD (black open: pre-induction; blue closed: post-induction; n = 8, p = 0.0006, Two-way RM ANOVA). Insets show representative traces of depolarization-induced AP train. Scale: 200 ms (horizontal) and 20 mV (vertical). (E) Bar graphs showing the changes in PF-evoked spike count between conditioned PF and unconditioned PF (grey left: pre-induction at conditioned PF; blue: post-induction at conditioned PF; grey right: pre-induction at unconditioned PF; purple: post-induction at unconditioned PF). There was significant changes in PF-evoked spike count when conditioned PF was stimulated (spike count: pre-induction = 15.38 ± 1.34; post-induction = 7.13 ± 0.64, n = 8, p = 0.008) while spike count was not changed when unconditioned PF was stimulated (spike count: pre-induction = 18.13 ± 3.00; post-induction = 15 ± 1.43, n = 8, p = 0.13). Insets show representative traces of PF-evoked spikes, elicited by stimulating 20 times of PF with 20 Hz. Scale: 250 ms (horizontal) and 20 mV (vertical). (F) Box and whisker plots showing the PF-evoked spike count from conditioned PF (blue) and unconditioned PF (black). Changes in PF-evoked spike output was prominent at the conditioned PF compared to unconditioned PF (∆spike count in conditioned PF = −52.28 ± 4.35% vs. unconditioned PF = −11.71 ± 5.97, p = 0.0006, Mann-whitney test). For statistics, Mann-whitney test was used for C (right) and F and Wilcoxon test was used for comparison of paired data set in E. Two-way RM ANOVA was used for D and post hoc Tukey’s test was used for within group (time) comparison. Error bar indicates SEM. n.s. denotes ‘not significant’, **p < 0.01, ***p < 0.001. * in panel D indicated statistical difference between each time point and significances was tested by post-*hoc* tukey test of Two-way RM ANOVA.

### Sufficient changes in spiking output requires both synaptic and intrinsic plasticity

Would the spiking output of PCs reflect either PF-PC LTD or LTD-IE? To clarify this, we pharmacologically inhibited PF-PC LTD without excitability changes by applying the first small-molecule inhibitor (FSC231, 50 µM) which prevents internalization of α-amino-3-hydroxy-5-methyl-4-isoxazolepropionic acid (AMPA) receptors (Thorsen et al., 2010). The FSC231 effectively inhibited PF-PC LTD but not LTD-IE (figure 3A – D). PF-evoked spiking activity was found to be decreased from both DMSO-treated control and FSC231-treated slices after the induction of PF-PC LTD, However, the variation of spike count (∆spike count) in the DMSO-control group was more prominent compared to that of FSC231-treated group (−65.15 ± 5.70% vs. −31.76 ± 11.85%, DMSO and FSC231, respectively, p = 0.04, t-test; figure 3E and F). Next, we further tested if excitability changes *per se* would be reflected in spiking output. We used a transgenic mice model, the PC-specific stromal interaction molecule 1 knockout mice (STIM1^PKO^), previously reported the impairment of intrinsic plasticity without the deficit of synaptic plasticity (Jang et al., 2019; Ryu et al., 2017). Consistent with a previous study, the synaptic plasticity was comparable between genotypes but the STIM1^PKO^ exhibited a deficit of LTD-IE (figure 3G – J). As the data presented above showed, PF-evoked spike counts decreased following LTD induction from both genotypes (figure 3K). The extent of changes in spike output after LTD induction in STIM1^PKO^, however, was less than the value from wild-type littermates (STIM1^WT^) (−64.17 ± 6.48% vs. −26.17 ± 4.18%, STIM1^WT^ and STIM1^PKO^, respectively, p = 0.004, Mann-Whitney test; figure 3K and L). Altogether, the activity-dependent modulation of neuronal output requires synergistic coordination of synaptic and neuronal excitability change.

**Figure 3.**
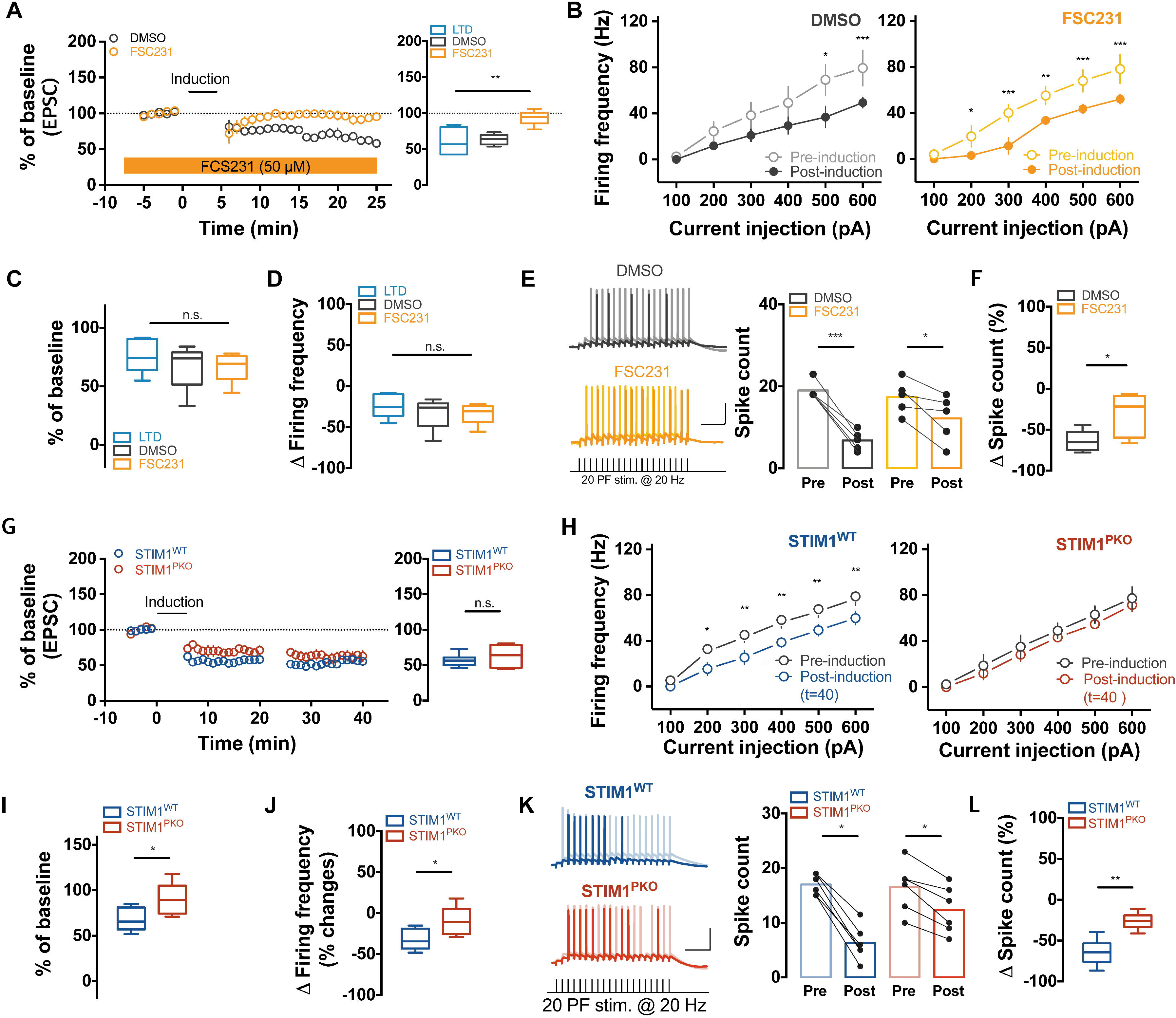
Activity-dependent modulation of PC spike output required synergies between synaptic and intrinsic plasticity. (A) Plots (left) showing the normalized eEPSC before and after LTD induction in a presence of PICK1 inhibitor, FSC231 (50 µM, orange, n = 5) and DMSO control (1:1000, black, n = 5) and summarizing box and whisker plots (right) of changes in eEPSC. LTD (blue) indicated the value shown in the LTD_ISI=120_ of figure 1C. Compared to LTD, FSC231 prevented induction of PF-LTD (% of baseline: DMSO control = 63.33 ± 3.33%, p = 0.86; FSC231 = 93.60 ± 4.61%, p = 0.001, One-way ANOVA post-*hoc* tukey test compared to LTD). (B) Plots showing frequency – current (F/I) curve of DMSO control and FSC231-treated group corresponding to time after induction (grey open: DMSO pre-induction; black closed: DMSO post-induction, n =5, F(5, 20) = 4.64, p = 0.006; orange open: FSC231 pre-induction; orange closed: FSC231 post-induction, n = 5, F(1, 4) = 8.88, p = 0.04, Two-way RM ANOVA). LTD-IE was exhibited in both DMSO- and FSC231-treated groups. Insets show representative traces of depolarization-induced AP train. Scale: 200 ms (horizontal) and 20 mV (vertical). (C - D) Box and whisker plots showing the changes in excitability after LTD induction throughout the groups including LTD (blue), DMSO-(black) and FSC231-treated groups (orange). There were no significant differences of excitability change between groups [(C): F(2, 12) = 0.61, p = 0.56; (D): F(2, 12) = 0.61, p = 0.56, One-way ANOVA]. (E) Bar graphs showing the changes in PF-evoked spike count between before and after induction in a presence of FSC231 (light orange: pre-induction; orange: post-induction) and DMSO (grey: pre-induction; black: post-induction). The PF-evoked spike count before and after LTD induction was significantly reduced in both groups (spike count: DMSO pre-induction = 19.00 ± 1.00; post-induction = 6.80 ± 1.07, n = 5, p = 0.0006, FSC231 pre-induction = 17.4 ± 1.86; post-induction = 12.2 ± 2.538, n = 5, p = 0.039, paired t-test). Insets show representative traces of PF-evoked spikes, elicited by stimulating 20 times of PF with 20 Hz. Scale: 250 ms (horizontal) and 20 mV (vertical). (F) Box and whisker plots showing the PF-evoked spike count from DMSO (black) and FSC231-treated group (orange). The PF-evoked spike count showed less decrease in the FSC231 treated group compared to DMSO control (∆spike count: DMSO = −65.15 ± 5.70% vs. FSC231 = −31.76 ± 11.85%, p = 0.04, t-test). (G) Plots (left) showing the normalized eEPSC before and after LTD induction from STIM1^WT^ (blue, n = 6) and STIM1^PKO^ (red, n = 6) and summarizing box and whisker plots (right) of changes in eEPSC. The changes in eEPSC was comparable between genotypes (% of baseline: STIM1^WT^ = 56.71 ± 3.68% vs. STIM1^PKO^ = 62.92 ± 6.47%, p = 0.82, Mann-Whitney test). (H) Plots showing frequency – current (F/I) curve of STIM1^WT^ (left) and STIM1^PKO^ (right) corresponding to time after induction. LTD-IE was impaired in STIM1^PKO^ while LTD-IE was intact in wildtype littermates [STIM1^WT^: black = pre-induction; blue = post-induction, n = 6, F (5, 25) = 78, p < 0.001; STIM1^PKO^: black = pre-induction; red = post-induction, n = 6, F(1, 5) = 1.42, p = 0.29, Two-way RM ANOVA]. Insets show representative traces of depolarization-induced AP train. Scale: 200 ms (horizontal) and 20 mV (vertical). (I - J) Box and whisker plots showing the changes in excitability after LTD induction throughout the groups between genotypes. There were significant differences of excitability change between genotypes [(I): p = 0.03; (J): p = 0.03]. (K) Bar graphs showing the changes in PF-evoked spike count between before and after induction from STIM1^WT^ (light blue: pre-induction; blue: post-induction) and STIM1^PKO^ (light red: pre-induction; red: post-induction). The PF-evoked spike count before and after LTD induction was significantly reduced in both groups (spike count: STIM1^WT^ pre-induction = 17.00 ± 0.77; post-induction = 6.25 ± 1.13, n = 6, p = 0.03, STIM1^PKO^ pre-induction = 16.5 ± 1.84; post-induction = 12.33 ± 1.76, n = 6, p = 0.03, Wilcoxon test). Insets show representative traces of PF-evoked spikes, elicited by stimulating 20 times of PF with 20 Hz. Scale: 250 ms (horizontal) and 20 mV (vertical). (L) Box and whisker plots showing the PF-evoked spike count from STIM1^WT^ (blue) and STIM1^PKO^ (red). The PF-evoked spike count shored less decrease in the STIM1^PKO^ compared to STIM1^WT^ (∆spike count: STIM1^WT^ = −64.17 ± 6.477% vs. STIM1^PKO^ = −26.17 ± 4.18%, p = 0.004, Mann-Whitney test). For statistics, One-way ANOVA test was used for A, C and D and post-hoc tukey test was used for different individual group comparison and two-way RM ANOVA was used for B and H and post hoc Tukey’s test was used for different time group comparison. Wilcoxon test was used for paired data set of E and K and Mann-Whitney test was used for F, G, I, J and L. Error bar indicates SEM. n.s. denotes ‘not significant’; *P < 0.05, **p < 0.01, ***p < 0.001. * in panel B and H indicated statistical difference between each time point and significances was tested by post-hoc tukey test of Two-way RM ANOVA.

In summary, our recordings from floccular PCs revealed that the postsynaptic spiking output reflects the synaptic plasticity by linking intrinsic plasticity. Since the intrinsic plasticity is modulated in the same polarity to the synaptic plasticity, the changes in the excitability has been assumed to amplify the changes in synaptic efficacy (Belmeguenai et al., 2010; Shim et al., 2017). In this work, we further elucidate the several physiological features of intrinsic plasticity in cerebellar PCs. First, the intrinsic plasticity in the floccular PCs follows the distinct timing rules governing PF-PC LTD by sharing the intracellular signaling cascade such as mGluR1-PKC and CaMKII. Second, the intrinsic plasticity of PCs is formed at the conditioned dendritic branches. These results provide an insight into synergistic coincidence of synaptic and intrinsic plasticity.

Multiple lines of evidence have demonstrated the heterogeneity of individual dendritic branches (Fu et al., 2012; Govindarajan et al., 2011; Ohtsuki et al., 2012; Zang et al., 2018). Thus, each dendritic branches may provide computational compartmentation enabling PCs to efficiently maximize the information storage capacity and actively integrate synaptic inputs (Ohtsuki et al., 2012; Zang et al., 2018). In fact, the branch specificity has been also implicated in clustered plasticity model, describing that the adjacent synaptic sites form functional clustering along dendritic branches in that similar information is preferentially processed in the clustered synapses (Fu et al., 2012; Govindarajan et al., 2011). Notably, spatiotemporal patterns of synaptic inputs encoding sensory information to the cerebellar PC dendrites are found to be activated in clusters (Wilms and Häusser, 2015). In this scenario, a spatiotemporal patterns of sensory information may form both synaptic and intrinsic plasticity at the specific branches of PC dendrite corresponding to similar sensory information. Hence, the PC output may reflect the synaptic inputs from only the conditioned dendritic sites whereas synaptic inputs from unconditioned synapses might be ignored. Taken together, the way to process the information in the cerebellar cortex is highly structured and clustered, therefore, localized formation of synaptic and intrinsic plasticity may play a significant role in information storage and furthermore modify the behavioral outcomes.

## Acknowledgement

We thank Yong-Seok Lee in Seoul National University, Jennifer Raymond in Stanford University and Geehoon Chung in Kyunghee University for providing scientific advices and Misun Mun, Jewoo Seo and Alex Fanning for proofreading this manuscript.

