## Supplemental information for "Synaptic and intrinsic plasticity coordinate spike output in cerebellar Purkinje cells"

Material and Methods

Supplemental Figure 1

**Material and Methods**

1. *Animals and Slice preparation*

All animal use was in accordance with protocols approved by the Institution’s Animal Care and Use Committee of Seoul National University College of Medicine. Cerebellar coronal slices were dissected into 250 μm by vibratome (Leica, VT1200) from anaesthetized 4- 6 weeks old male C57BL/6 mice in ice-cold standard artificial cerebrospinal fluid (aCSF) contained with the following (in mM): 125 NaCl, 2.5 KCl, 1 MgCl_2_, 2 CaCl_2_, 1.25 NaH_2_PO_4_, 26 NaHCO_3_, 10 glucose bubbled with 95% O2 and 5% CO2. For recovery, slices were incubated at 32 ˚C for 30 minutes and further 1 hour at room temperature.

2. *Electrophysiology*

Slices were put onto a submerged recording chamber on the stage of Olympus microscope (BX50WI, Japan) and perfused with aCSF (composition is described above), and kept in place with a nylon-strung platinum anchor. All recordings were performed using multiclamp 700B patch-clamp amplifier (Axon Instruments) with a sampling frequency of 20 kHz and signals were filtered at 2 kHz under a presence of inhibitory synaptic transmission inhibitor (picrotoxin, 100 μM) to isolate the excitatory synaptic transmission. Patch pipettes (3-4 MΩ) were borosilicate glass and filled with internal solution containing (in mM), 9 KCl, 10 KOH, 120 K-gluconate, 3.48 MgCl_2_, 10 HEPES, 4 NaCl, 4 Na_2_ATP, 0.4 Na_3_GTP, and 17.5 sucrose, pH adjusted to 7.25. The membrane potential was held on -70 mV in voltage-clamp (VC) and current-clamp mode (CC). Recordings were discarded if the series resistance (R_s_) varied by > 15% and the injection current for the holding potential exceeded 600 pA. PFs in the molecular layer and CF in the granule cell layer were stimulated by an ACSF-filled electrode. We used three different protocols to induce the PF-PC LTD in CC mode for 5 min: 1) tetanizing of PF and CF simultaneously (1 Hz, 300 times for 5 min, PF-LTD_ISI=0_; figure 1B left), 2) tetanizing of PF followed by single CF stimulation with the stimulus interval of 120 ms (1Hz, 300 times for 5 min, PF-LTD_ISI=120_; middle), 3) 7 times of burst stimulation of PF with 100 Hz followed by single CF stimulation with the stimulus interval of 150 ms (30 times and sweep interval 10 s for 5 min; PF-LTD_ISI=150burst_; right) (Shim et al., 2017; Suvrathan et al., 2016). To evaluate the PCs excitability, a series of current steps of 500 ms duration ranging from +100 pA to +500 pA with increments of 100 pA with a step interval of 4.5 s from a membrane potential of -70 mV was injected in CC mode.

3. *Data acquisition and analysis*

All data were acquired by Clampex software (Molecular Devices) and analyzed by IgorPro 8.1 (Wavemetrics). We evaluated the LTD by normalizing to the average of baseline PF-EPSC, and recording was excluded if the baseline current or the number of evoked spikes varied by > 20 %. R_S_ was calculated by fitting a single exponentials to the voltage responses of the test pulse (-5 mV).

Data were presented as mean ± SEM and statistical evaluations were performed using normality test, equal variant test, independent *t*-test, Mann-Whiteny test, One-way repetitive measured (RM) ANOVA and Two-way RM ANOVA with post *hoc* tukey test and paired t-test and Wilcoxon test were used for comparing paired data set by SigmaPlot 12.0 (Systat Software)and prism 7.0 (Graphpad) software. Sample size was approved by power analysis using the G*power 3.1.9.2.

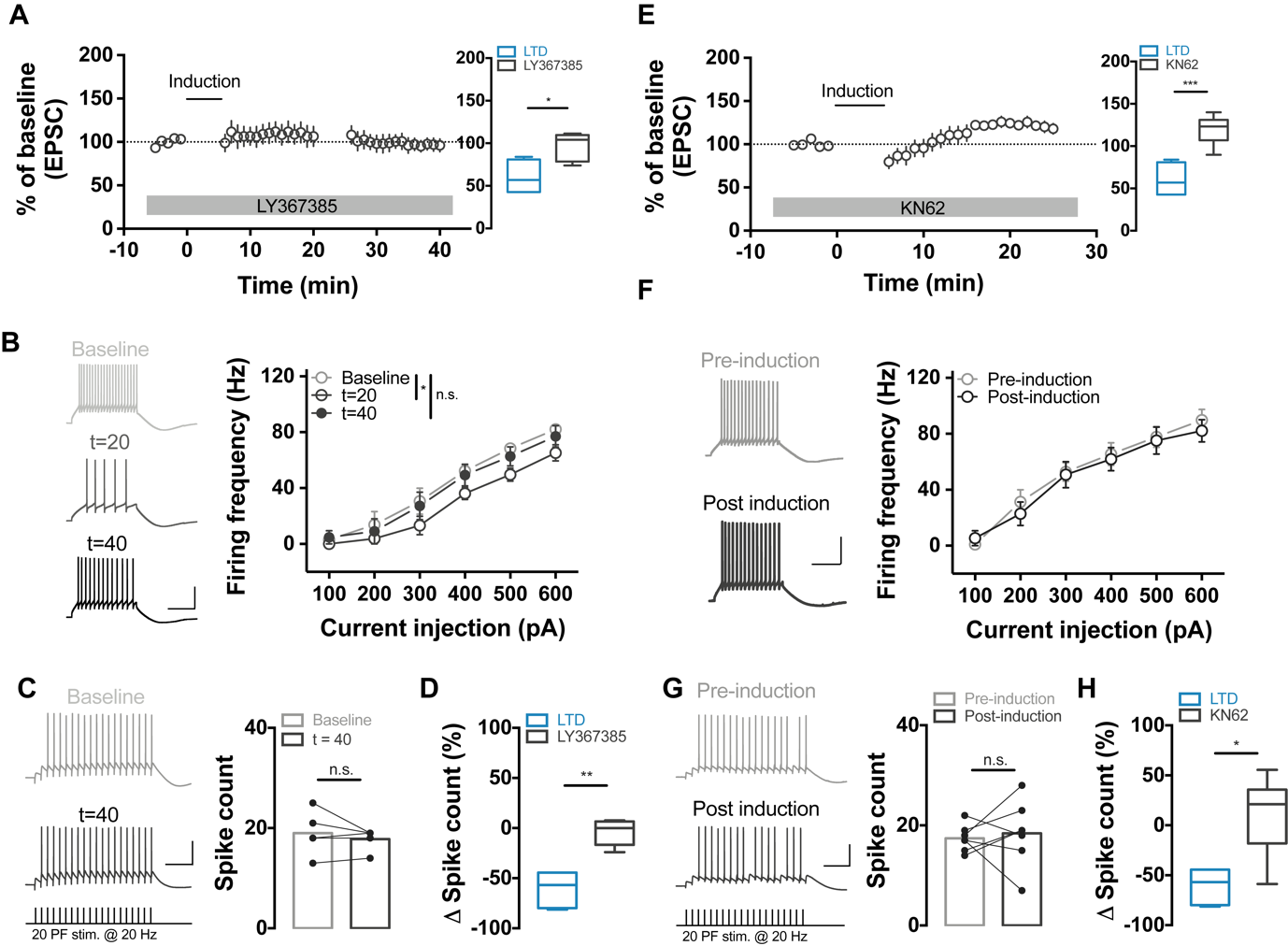

**Supplemental figure 1 (related to figure 1). The LTD-IE is dependent of mGluR1 signaling.**

(A) Plots (left) showing the normalized eEPSC before and after LTD induction in a presence of mGluR1 inhibitor, LY367385 (100 µM) and summarizing box and whisker plots (right) of changes in eEPSC corresponding to ISI_120_ (blue, value shown in fig. 14B, n = 7) and LY36738-treated group (black, n = 5). Compared to ISI_120_, LY367385 prevented induction of PF-LTD (p = 0.02, Mann-whitney test).
(B) Plots showing frequency – current (F/I) curve of LY367385-treated group corresponding to time after induction (grey: baseline; black open: t = 20; black closed: t = 40, n = 4, p = 0.10, Two-way RM ANOVA). The statistical difference was shown only at t = 20 compared to baseline and there was no significance between baseline and t = 40 (baseline vs. t = 20: p = 0.001; baseline vs. t = 40: p = 0.18, post-*hoc* tukey test). Insets show representative traces of depolarization-induced AP train. Scale: 200 ms (horizontal) and 20 mV (vertical).
(C) Bar graphs showing the changes in PF-evoked spike count between before and after induction in a presence of LY367385 (grey: baseline; black: t = 40). There was no significant changes in PF-evoked spike count before and after LTD induction (PF-evoked spike count: baseline = 19 ± 1.98 vs. t = 40 = 17.8 ± 0.97, p = 0.63, Wilcoxon test). Insets show representative traces of PF-evoked spikes, elicited by stimulating 20 times of PF with 20 Hz. Scale: 250 ms (horizontal) and 20 mV (vertical).
(D) Box and whisker plots showing the PF-evoked spike count from ISI_120_ (blue) and LY367385-treated group (grey). Inhibition of mGluR1 prevented the changes in PF-evoked spike output (∆spike count in LY367385- treated group = -4.06 ±5.81%, comparison with the value of ISI_120_, p = 0.01, Mann-whitney test).
(E) Plots (left) showing the normalized eEPSC before and after LTD induction in a presence of CaMKII inhibitor, KN62 (3 µM) and summarizing box and whisker plots (right) of changes in eEPSC corresponding to ISI_120_ (blue, value shown in fig. 14B, n = 7) and KN62-treated group (black, n = 7). Compared to ISI_120_, KN62 prevented induction of PF-LTD (p = 0.0003, Mann-whitney test).
(F) Plots showing frequency – current (F/I) curve of LY367385-treated group corresponding to time after induction (grey: pre-induction; black: post-induction, n = 5). Insets show representative traces of depolarization-induced AP train. Scale: 200 ms (horizontal) and 20 mV (vertical).
(G) Bar graphs showing the changes in PF-evoked spike count between before and after induction in a presence of KN62 (grey: pre-induction; black: post-induction). There was no significant changes in PF-evoked spike count before and after LTD induction (PF-evoked spike count: pre-induction = 17.43 ± 0.10 vs. post-induction = 18.43 ± 2.47, p = 0.63, Wilcoxon test). Insets show representative traces of PF-evoked spikes, elicited by stimulating 20 times of PF with 20 Hz. Scale: 250 ms (horizontal) and 20 mV (vertical).
(H) Box and whisker plots showing the PF-evoked spike count from ISI_120_ (blue) and KN62-treated group (grey). Inhibition of CaMKII prevented the changes in PF-evoked spike output (∆spike count in KN62- treated group = 7.17 ± 17.71%, comparison with the value of ISI_120_, p = 0.02, Mann-whitney test).

For statistics, Mann-whitney test was used for A (right), D, E (right) and H and Wilcoxon test was used for comparison of paired data set in C and G. Two-way RM ANOVA was used for B and post hoc Tukey’s test was used for different time group comparison. Error bar indicates SEM. n.s. denotes ‘not significant’; *P < 0.05, **p < 0.01, ***p < 0.001. * in panel B indicated statistical difference between each time point and significances was tested by post-*hoc* tukey test of Two-way RM ANOVA.
